# Novel mouse reporter models for the detection of genome editing events in vivo

**DOI:** 10.64898/2026.04.29.721708

**Authors:** Kathy J. Snow, Ethan Saville, Caleb Heffner, Yaned Gaitan, Jeffrey Duryea, Tiffany L. Davis, Lawrence Bechtel, Seth Hannigan, Benjamin E. Low, Jana Rossius, Tu Dang, Katarina Kulhankova, Anna X. Cheng, Michael V. Wiles, Wolfgang Wurst, Paul B. McCray, David Guay, Cathleen M. Lutz, David E. Bergstrom, Ralf Kühn, Stephen A. Murray

**Affiliations:** The Jackson Laboratory, Bar Harbor, ME, USA, 04609; Max-Delbrück-Center for Molecular Medicine in the Helmholtz Association (MDC), Berlin, Germany; Department of Pediatrics, University of Iowa, Iowa City, IA, USA; Feldan Therapeutics, Quebec, Qc, Canada; Department of Stem Cell Research, Helmholtz Zentrum München, Neuherberg, Germany

**Keywords:** gene editing, base editing, reporter mice, CRISPR, rare diseases

## Abstract

With the expansion of therapeutic gene editing technology, small animal models provide essential platforms to evaluate the function of these new approaches *in vivo*. As part of the Somatic Cell Genome Editing (SCGE) Consortium, we developed next-generation murine reporters that overcome current model limitations and broaden detectable *in vivo* editing outcomes. These include two mouse models built on the “traffic light” reporter concept. This system enables fluorescent detection of both gene repair (green) and CRISPR-generated indels (red) events following editing by a single guide and either dsDNA or single-stranded oligonucleotide donor. We also generated a third reporter model that efficiently detects A-base editor activity. Reporters were validated in cultured embryos, via germline editing, and through activation *in vivo* by AAV transduction or direct ribonucleoprotein delivery. Together, these new models provide a valuable resource for improved detection of genome editing events *in vivo*.

## Introduction

Programable nucleases such as CRISPR/Cas9 have revolutionized our ability to manipulate the genome. This has simplified the process of creating genetically modified cells and model organisms, decreasing the time required and reducing the associated costs. Moreover, the technology has greatly expanded the range of species that can be effectively engineered, enabling experimentation not previously possible.

The potential for therapeutic uses of CRISPR/Cas9 are particularly exciting, including both *ex vivo* and *in vivo* editing of somatic cells ^1^. These approaches promise to generate robust and durable modifications of the genome after a single treatment and several clinical trials have demonstrated remarkable efficacy including the recent approval of the *ex vivo* therapy Casgevy for the treatment of Sickle Cell Disease (Vertex Pharmaceuticals) and the remarkable N-of-1 treatment of an infant with severe carbamoyl phosphate synthetase 1 (CPS1) deficiency ^2–5^. Despite these promising initial results several challenges remain before the potential of therapeutic genome editing can be fully realized.

Delivery of genome editing cargo to the relevant cell population in an efficient and specific manner remains a major challenge for most tissues outside of the liver. Viral vectors, including recombinant adeno-associated virus (rAAV) have been used to great effect for gene replacement applications ^6,7^, but their clinical utility for genome editing has not been demonstrated. Similarly, lipid nanoparticles (LNPs) and modified ribonucleoproteins (RNPs) show great promise without the potential risk of long-term expression of an editor ^4^, but much work remains to develop delivery platforms that both efficiently and specifically target the cell and tissue type of interest. Given the complexity of factors that impact the distribution, bioavailability and pharmacokinetics of diverse delivery technologies, the use of small animals to evaluate genome editor delivery specificity along with editing efficiency will be a critical step in advancing new delivery platforms to the clinic.

Reporter mouse models allow for the simple evaluation of genetic manipulation at single-cell resolution *in situ* through the activation of a fluorescent or other visible marker. These tools have been widely used to both evaluate the specificity of cre recombinase driver strains and for lineage tracing experiments *in vivo* ^8–11^. An ideal reporter has several key characteristics, including low leakiness, insertion at a chromatin-accessible locus, robust activation of a detectable signal in a single cell following allele modification, durability of the signal over time, and potential for activation in all cells and tissues, ideally at uniform levels. Current cre reporter strains, such as Ai9 or Ai14 (^9^, JAX#007909, #007914), using a ubiquitous CAGGS promoter driving a bright fluorescent protein targeted to the accessible ROSA26 locus meet most of these criteria.

Because of these advantages, cre reporter strains have been redeployed for monitoring of genome editing activity following delivery of editor cargo ^12–14^. Typically, guides are designed to flank the lox-stop-lox terminator cassette activating the reporter when the cassette is excised ^13^. Alternatively, a single guide can be targeted to the repeated terminator sequence activating the reporter when one or more are subsequently removed ^12^. Both strategies rely on excision of an intervening sequence following multiple double-stranded breaks that are repaired by non-homologous end-joining (NHEJ) of the flanking DNA. Because activation requires that multiple nuclease events occur simultaneously in cis and repair via removal of the intervening sequence, some nuclease activity is missed; e.g. single cuts repaired by NHEJ or multiple cuts repaired independently with retention of the terminator cassette. Additionally, repair of multiple DS breaks has been shown to result in complex rearrangements, further complicating interpretation of the reporter signal. These reporters are limited to detection of nuclease events and are unsuitable for applications using homology-directed repair following DS breaks or base/prime editors, technologies that are of growing interest for therapeutic applications ^1,15–18^ .

Due to the limitations of the current reporter options, an expanded toolbox is required to support the testing and validation of new delivery technologies with diverse editor types and usage configurations. In addition to the key characteristics of a useful reporter described above, additional desirable features specific to gene editing reporters include responsiveness to a single guide, detection of indel generation following repair of single cut, detection of homology-directed repair (HDR) events, detection of base editor activity, and the potential to report both HDR and NHEJ in the same allele.

Here, we report three novel mouse reporter models that fill these gaps and provide new and readily accessible tools for the research community to advance the development of new delivery technologies for therapeutic genome editing.

## Results

### Development and validation of the traffic light reporter (TLR-2) mouse model

We adopted a “traffic light” design to construct a reporter allele that 1) would be activated by a single guide-mediated nuclease event; and 2) could detect both NHEJ and HDR events in a single allele, if a suitable double-stranded DNA donor was simultaneously delivered. As shown in **Figure 1a**, the Traffic Light 2 (TLR-2) allele includes a mVenus fluorescent protein interrupted by an 108 bp DNA segment ^19^ that eliminates its fluorescent potential but can be repaired with a DNA donor. A red fluorescent TagRFP cassette is linked by a P2A peptide sequence out of frame with mVenus, and thus the protein is not expressed in the native allele configuration. Upon NHEJ-mediated repair of double-stranded breaks, any indel that results in a frameshift to the +3 frame (e.g. 2bp deletion or 1bp insertion) will permit in-frame translation of the TagRFP protein. The TLR-2 cassette is driven by a ubiquitous CAGGS promoter/enhancer and targeted to the ROSA26 safe harbor locus similar to cre reporter alleles (see

**Figure 1.**
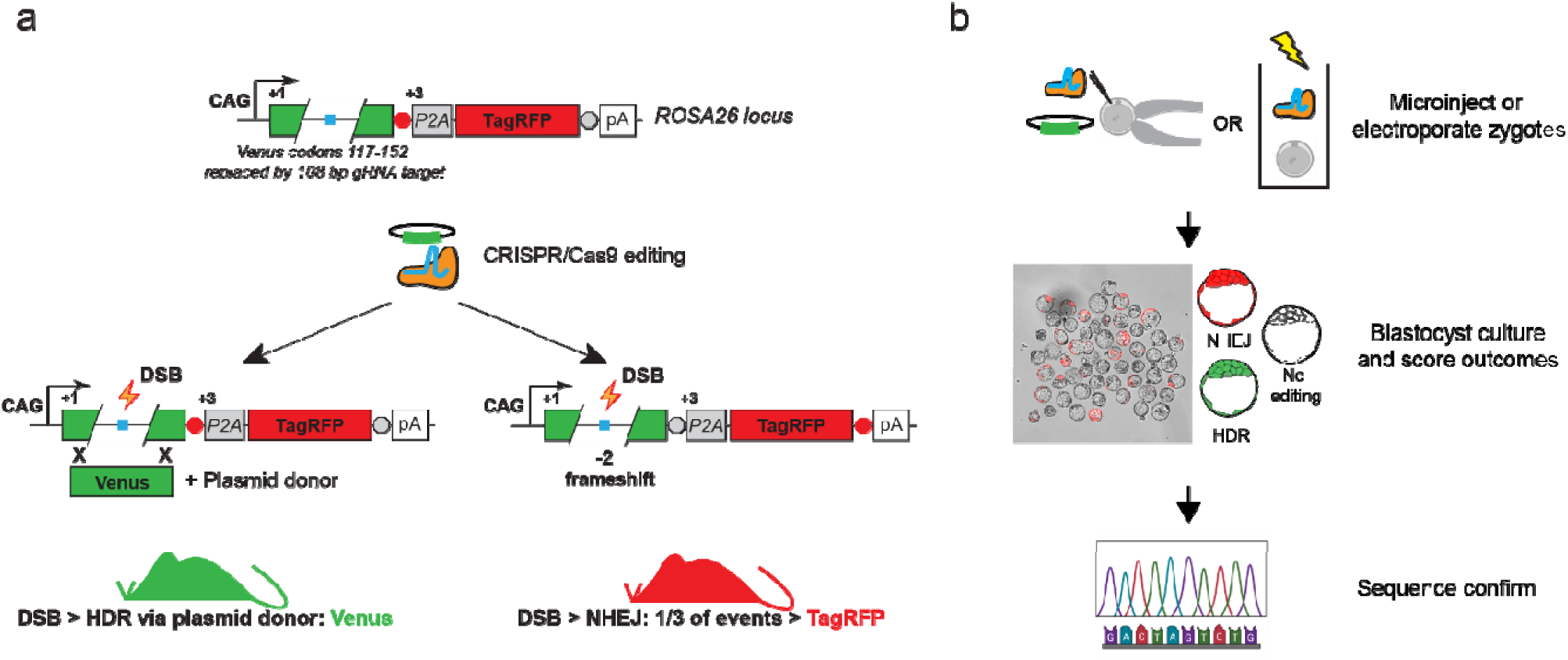
Traffic light reporter 2 (TLR-2): multifunctional allele for detection of both HDR and NHEJ editing events. **a)** Schematic of TLR-2 reporter allele structure and outcomes following editing. The allele includes two open reading frames encoding different fluorescent proteins; an upstream mVenus (green) in the +1 frame and downstream TagRFP (red) in the +3 frame linked by a P2A sequence. The mVenus cassette is interrupted by a 108 bp polylinker sequence that includes several guide target sequences and a stop codon (blue square). Thus in its native configuration, neither fluorescent protein is expressed. Following CRISPR/Cas9 editing, repair via NHEJ will result in expression of TagRFP if the frame shift results in a -2 deletion, or multiple thereof. Alternatively, HDR can be detected with the inclusion of a plasmid donor designed to repair the mVenus gap. **b)** Mouse embryo reporter validation workflow. Fertilized zygotes from TLR-2 reporter mice (typically male homozygous to WT female) are either microinjected or electroporated with Cas9 RNP with or without a plasmid donor for HDR. The embryos are then cultured to the blastocyst stage where the outcome can be scored by fluorescent imaging, and followed up by PCR-Sanger sequencing to confirm the identify of specific edits.

Materials and Methods for details on animal model generation). We developed a mouse embryo culture assay that would allow us to rapidly evaluate the relative efficiency of any guide sequence for editing activity and activation of TagRFP **(Figure 1b)**. Mouse zygotes harboring the TLR-2 allele (heterozygous) are harvested at 0.5 dpc, electroporated with Cas9 RNPs complexed with the specific guide of interest, and cultured to the blastocyst stage where they were visualized and scored for TagRFP signal. Following imaging, the blastocysts are collected and individually genotyped to determine the level of editing and to determine the specific indel outcomes.

To validate the functionality of the TLR-2 allele, we designed and tested a panel of guides to assess their ability to activate the TagRFP reporter. Several guide sequences for SpyCas9 and SauCas9 in the linker sequence or upstream portion of mVenus 5’ to a stop codon can be used to activate the TagRFP allele for monitoring nuclease activity *in vivo*. Guide locations relative to the mVenus coding and linker sequence in the TLR-2 are shown in **Figure 2a** and sequences provided in **Supplementary Table S1**. Of note, the R26-1 guide cut site is 3’ to the stop codon and thus is predicted to be a poor guide for detecting NHEJ events but could be useful for driving HDR. Three of the upstream guides (R26-52, R26-59, and R26-69) were selected using inDelphi ^20^ (https://indelphi.giffordlab.mit.edu/) for repair events following SpyCas9 induced dsDNA breaks that are predicted to result in an optimal shift into the +3 frame which is required to activate TagRFP. As shown in **Figure 2b**, we observed between 10 and 60% activation by fluorescence and 73% and 93% by editing using SpyCas9 guide RNP complexes. As noted above, the large discrepancy between the high editing rate and low fluorescent activation for guide R26-1 is attributable to the position of the cut 3’ of the stop codon in the linker sequence and the preference for +1 indels following repair. By contrast, guide R26-52 showed both high levels of overall editing (83%) and the highest percentage of fluorescence of all guides tested, consistent with the inDelphi-predicted preference for +3 frame-shifting indels. Embryos homozygous for the reporter allele showed a higher rate of fluorescence activation, as expected, although the total rate was not additive (**Figure 2d and 2e**). We also tested the utility of the TLR-2 allele for monitoring of HDR using a microinjected double stranded plasmid donor in our blastocyst assay. As shown in **Supplementary Figure S1**, we were able to detect mVenus fluorescence in embryos, indicating successful HDR. This platform could be useful to test modifications of mouse model engineering protocols to improve efficiency of HDR.

**Figure 2.**
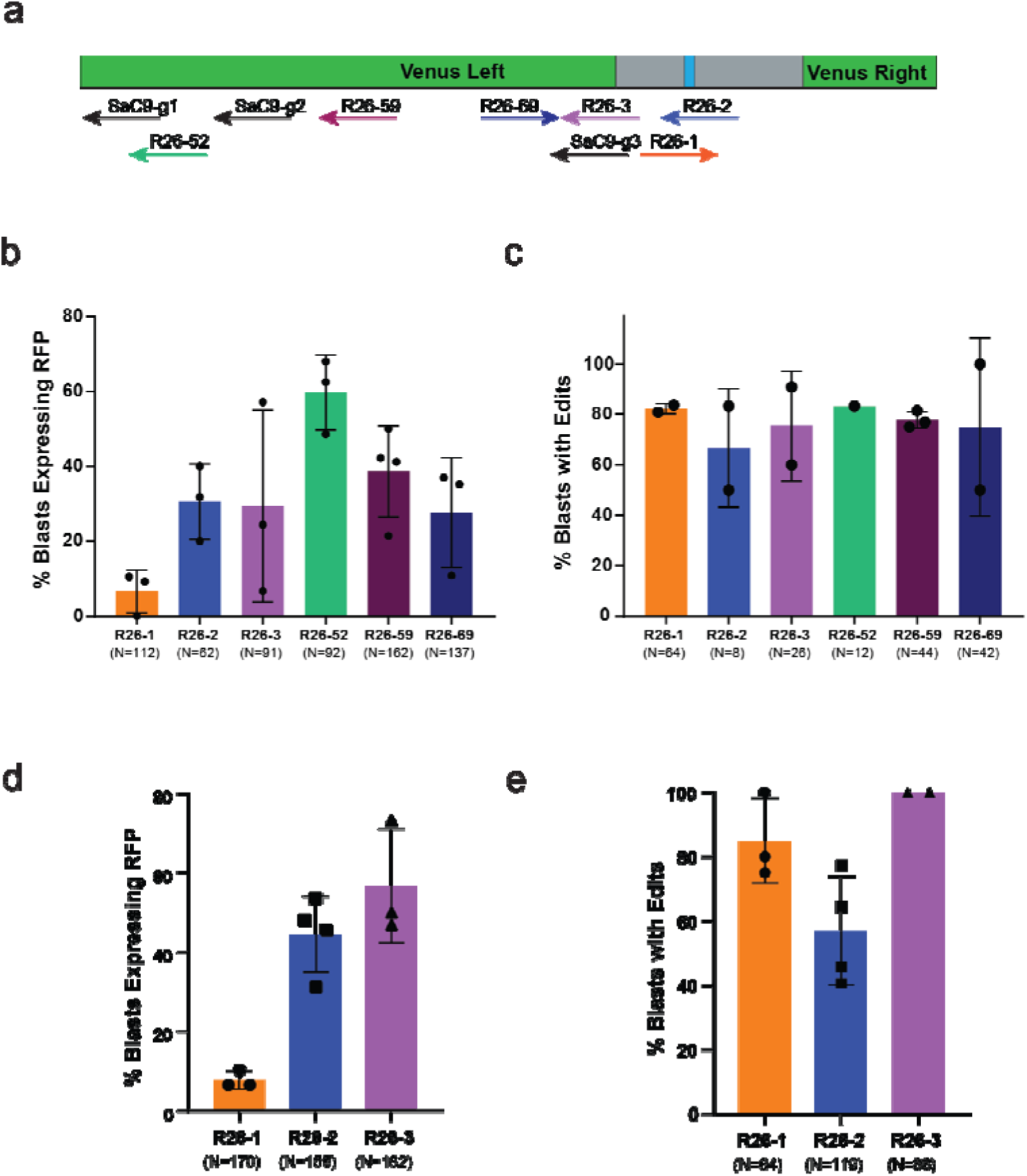
Evaluation of TLR-2 NHEJ detection efficiency in mouse embryos. **a)** Position of guides tested for the TLR-2 allele relative to the 108bp polylinker (gray) and TGA stop codon (blue). **b)** Percentage of TLR-2 heterozygous blastocysts showing visible expression of TagRFP following editing. Mouse zygotes electroporated with Cas9 RNPs with the guides listed on the X axis were cultured to the blastocysts stage and scored for expression of TagRFP, with corresponding levels of overall editing presented in **(c)**. Of note, the low number of blasts showing TagRFP fluorescence relative to the editing rate for R26-1 reflects its cut position 3’ to the stop codon in the polylinker. **d)** Percentage of TLR-2 homozyogous blastocysts with visible expression of TagRFP following editing and corresponding editing percentages shown in **(e)**. Data are representative of at least two technical replicates per guide and plots are showing mean and standard deviation of replicates with total number of blasts assessed across all replicates below each bar.

Generalized utility of a reporter model requires that the reporter cassette is expressed in all cells and tissue when activated. Although the proven CAGGS promoter/enhancer in the ROSA26 locus is predicted to provide for ubiquitous expression, we sought to validate this by editing our reporter in the germline and confirming lines harboring a +3 frame edit showed widespread, ubiquitous expression at adequate levels for detection (**Figure 3a**). As shown in **Figure 3b**, we found that all tissue samples showed robust expression of the TagRFP allele following immunostaining with an anti-RFP antibody due to the sensitivity of TagRFP to our fixation procedure. We observed some variation in fluorescent intensity between tissues, for example liver versus lung or kidney, which could reflect differences in the strength of the CAGGS promotor and/or the overall cellular density of the tissue. Representative images for additional tissues examined are shown in **Supplementary Figure S2**. Critically, we found robust activation in the germline, with clear expression in the sperm and oocyte, which allows for evaluation of the risk of germline editing for novel delivery formulations.

**Figure 3.**
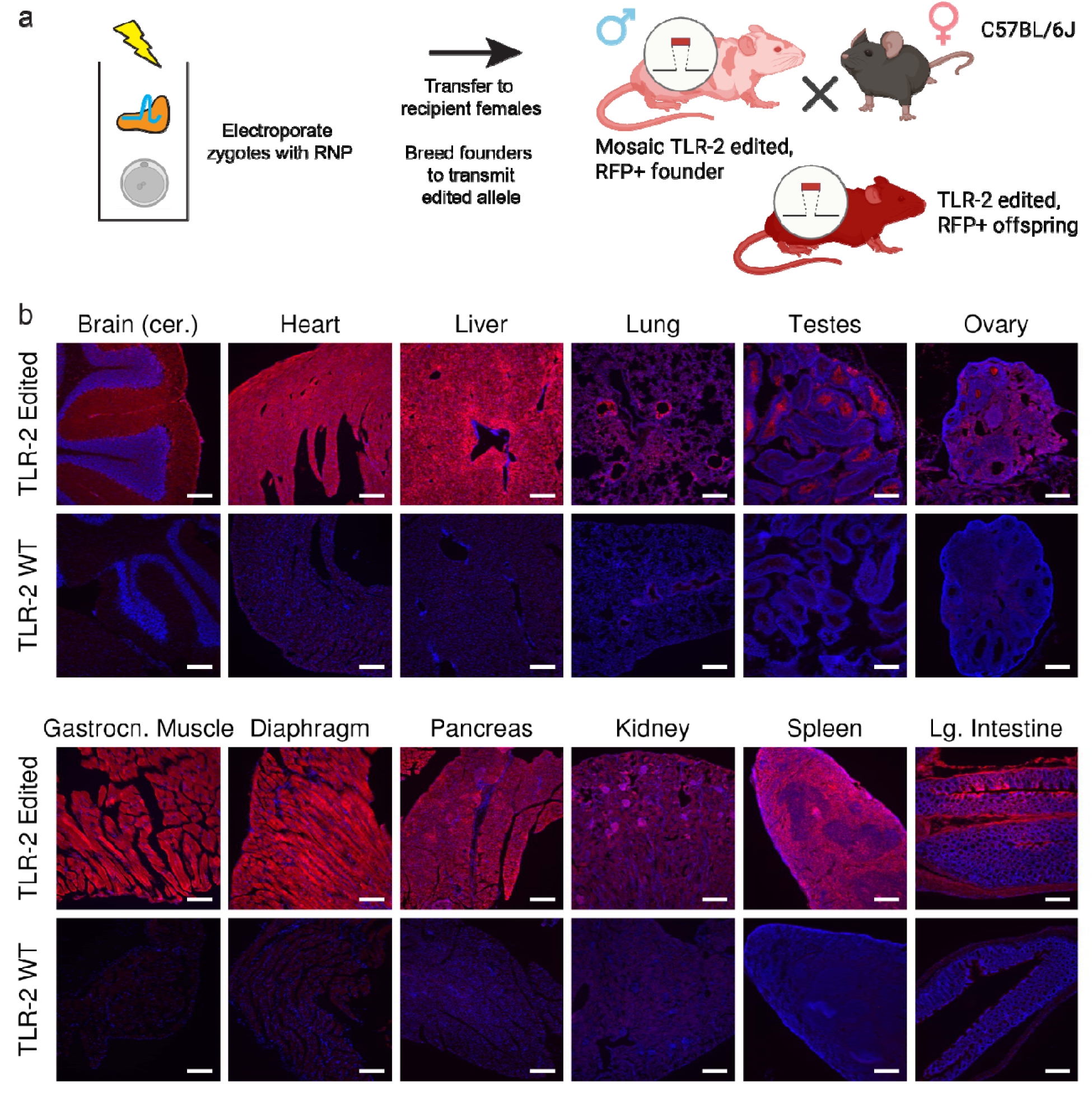
Validation of ubiquitous reporter expression following germline editing. **a)** To confirm the TLR-2 allele can be activated in all cells and tissues, we generated a germline model following targeting with Cas9 RNP and R26-52 guides. Founders harboring a -14 bp frameshift allele were bred to generate N1 mice. **b)** Imaging of 12 tissues showing widespread and consistent activation of TagRFP expression. Scale bar = 200 microns.

### *In vivo* rAAV delivery activates the TLR-2 reporter in transduced tissues

Finally, we sought to confirm the utility of our line for detecting only nuclease activity following *in vivo* delivery of an AAV-packaged gene editor. Since SauCas9 is often used in AAV-based delivery systems due to its reduced size, we searched the upstream mVenus sequence in the TLR-2 reporter for traditional SauCas9 guide and PAM target sites. We identified six possible guides and tested three (SaC9-g1, SaC9-g2, SaC9-g3) again using our embryo editing and blast culture assay (**Figure 4a,b**). Although two of the tested guides showed strong editing rates of >70% (**Figure 4b**), fluorescence activation was low versus the SpyCas9 guides (**Figure 4a**), as predicted by inDelphi for activating frameshifts, activation rates were sufficient for functional validation *in vivo*. Systemic (intravenous) delivery of AAV9-SauCas9-g3 virus (**Figure 4c**) led to robust editing in the liver as assessed by sequencing (**Figure 4d**) or immunofluorescence (**Figure 4e**), with detectable activation in the heart, and adrenal glands (**Supplementary Figure S3**). We observed greater apparent editing efficiency in males, consistent with previously reported sex effect ^21–23^ on AAV transduction in mice. Editing levels and frameshift proportions were roughly similar to what was seen in our embryo assay, demonstrating the utility of the embryo assay for predicting outcomes in adult animals, at least for the liver. As part of the SCGE program ^24^, the TLR-2 model was also deployed for our validation of a novel Focused Ultrasound (FUS) delivery system ^25^. An rAAV-SaCas9-guide delivered intravenously along with formulated microbubbles coupled with a tiled focused ultrasound array were able to edit specific sections of the mouse brain with very high efficiency (Figure 2G, ^25^). Taken together, these results demonstrate that the TLR-2 reporter is a useful model for assessment of *in vivo* genome editing delivery technologies.

**Figure 4.**
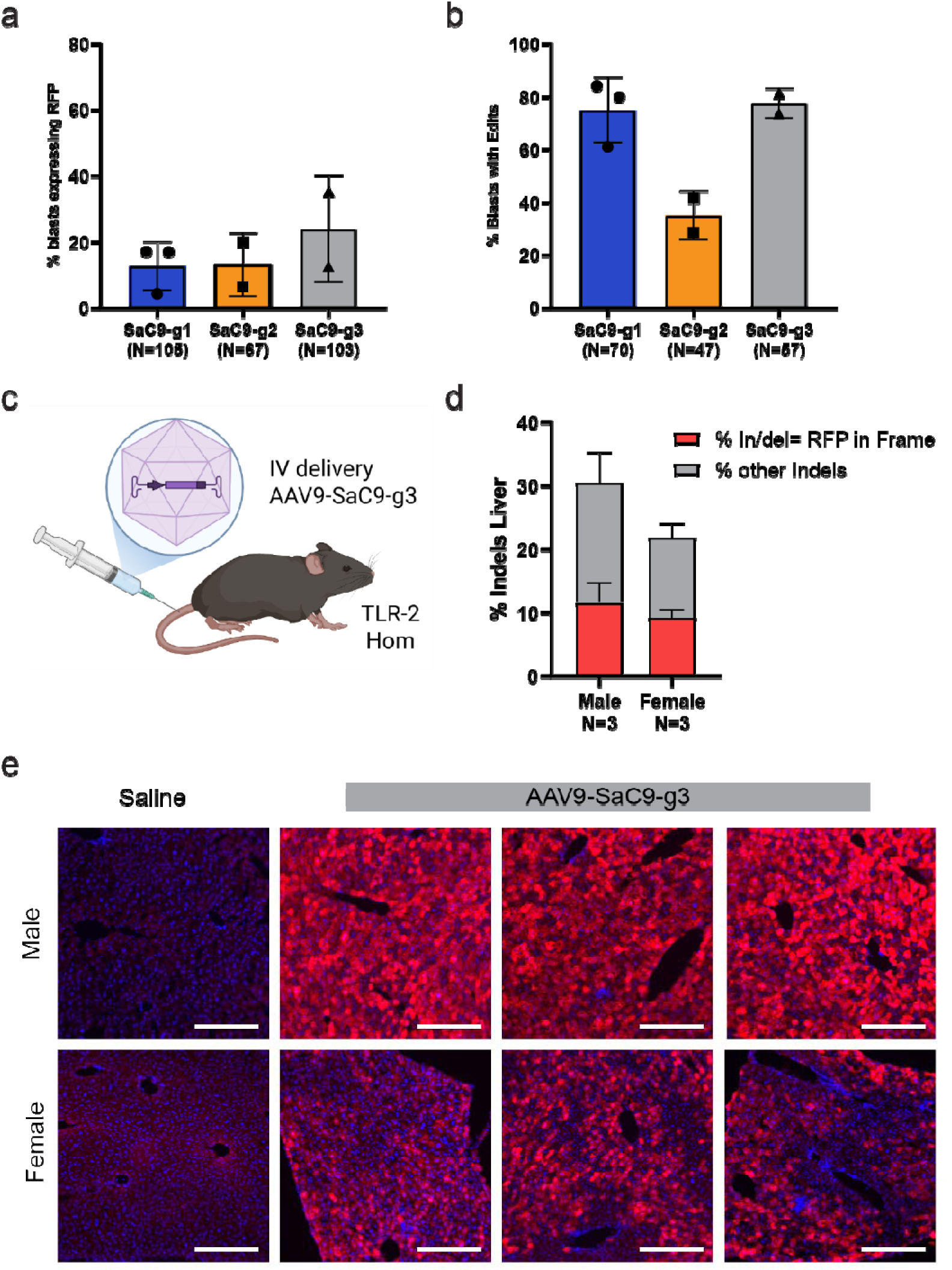
*In vivo* validation of TLR-2 reporter using AAV-mediated delivery of SauCas9. Embryo culture testing of SauCas9 guide efficiency for both fluorescence activation **(a)** and editing efficiency **(b)**. Plots are showing mean and standard deviation of replicates with total number of blasts assessed across all replicates below each bar. SaC9-g3 showed the best performance with ∼25% fluorescence and an 80% editing rate. **c)** AAV9 vectors encoding SauCas9 and guide SaC9-g3 were injected IV into 10 week old adult TLR-2 mice (1 x 10^11^vg/mouse) and tissues were harvested 4 weeks later (n=8, including controls). Genotyping of the liver **(d)** showed up to 30% editing of the TLR-2 allele of which 1/3 were in the TagRFP frame. Mean editing and Standard Deviation is shown for the AAV9-treated animals. **e)** Immunofluorescence of the liver of AAV9-treated male and female mice, showing substantial RFP activation versus saline control. Scale bar = 200 microns.

### Development and validation of the traffic light reporter 7 (TLR-7) mouse model for improved HDR detection

Although the TLR-2 model can be used to detect HDR events *in vivo* using a double stranded DNA donor, the large size of the linker inserted into the mVenus gene limits its utility for single-stranded oligonucleotide (ssODN)-mediated HDR events. To address this issue, we designed an additional traffic light-based reporter (TLR-7) that includes a smaller linker where 9 base pairs of the mVenus protein are replaced with a 30bp linker containing a single guide and PAM sequence target (**Figure 5a**). We also replaced the TagRFP with a Katushka2S fluorescent protein, which is red-shifted and has improved spectral qualities for live reporter imaging ^26^. We tested the TLR-7 reporter in our embryo assay and found that guides targeted to the linker sequence (TLR-7 g2_rev) produced indels at high efficiency (**Figure 5b**; 96%), with ∼35% of embryos showing red fluorescence indicating a frameshift into the +3 frame. To test the functionality for detecting ssODN-mediated repair, we included a 200bp repair oligo in the electroporation mix and found an average of 27.3% of edited embryos showed correct repair as indicated by mVenus fluorescence (**Figure 5c**). We confirmed this result by Sanger sequencing which showed a similar level of correct editing. As with TLR-2, we generated germline edited animals to confirm that the reporter protein is expressed in all cells and tissues (**Figure 5d**). Together, these data show that TLR-7 provides a useful tool for evaluation of both indel generation and ssODN-mediated HDR.

**Figure 5.**
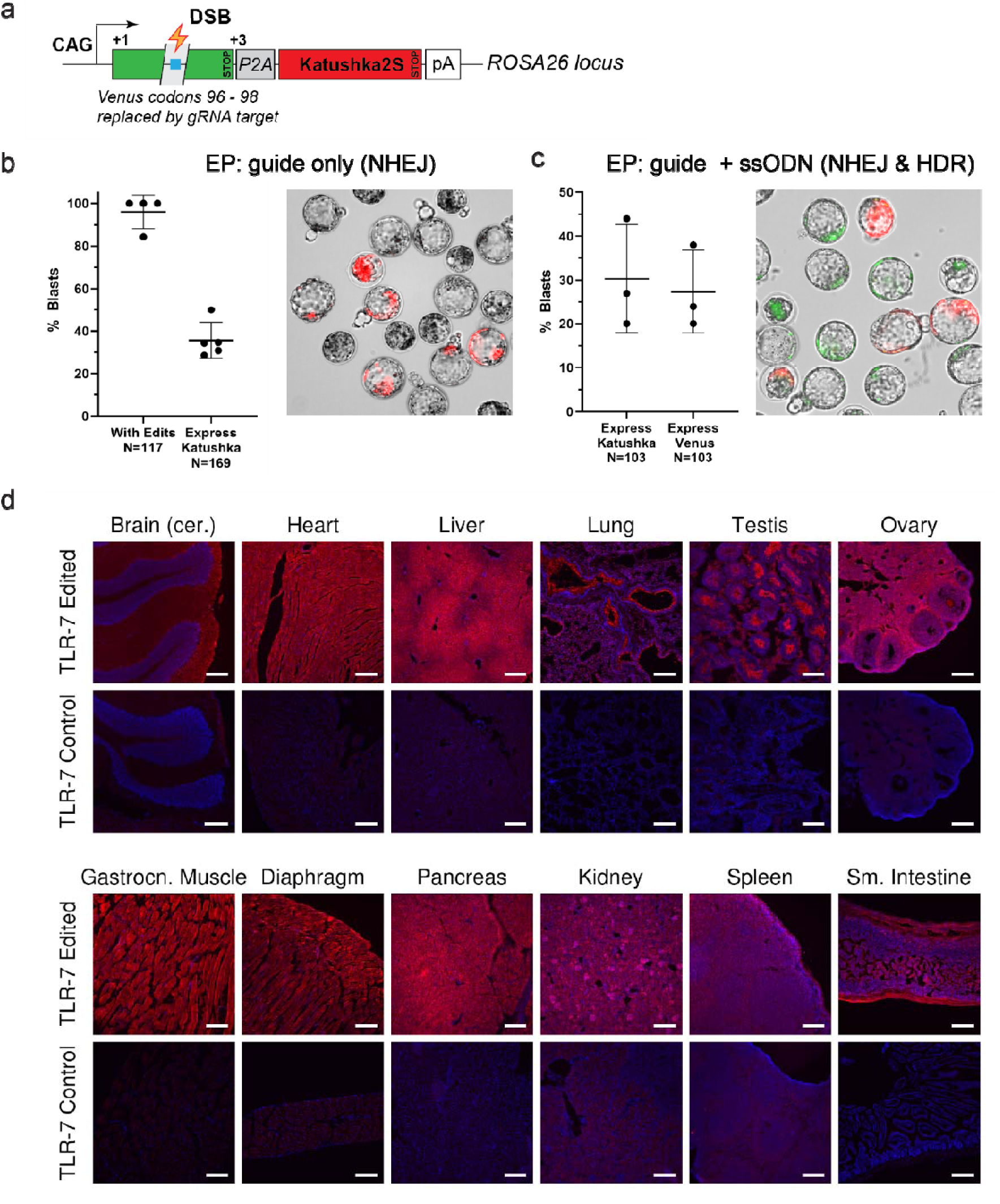
Single-strand oligonucleotide (ssODN) donor HDR compatibility in the TLR-7 reporter mode . **a)** Schematic of the TLR-7 allele, showing the shortened polylinker sequence (30 nt) as compared to TLR-2 including a stop codon (blue), that can be repaired with an ssODN. **b)** Mouse blastocyst validation for NHEJ repair shows high editing rates (>95%) with an expected 1/3 of edits resulting in a shift to the +3 frame to produce the red fluorescent signal. **c)** Validation of HDR functionality of the allele, showing ∼30% repair indicated by mVenus fluorescence and a similar level of NHEJ activation of Katushka2S. Some blastocysts showed mosaic activation of both cassettes. Plots are showing mean and standard deviation of replicates with total number of blasts assessed across all replicates below each bar. **d)** Germline indel generation (2 base pair deletion) demonstrates robust, widespread Katushka2S activation in all tissues. Scale bar = 200 microns.

### Generation of the gene editing reporter 10 (GER-10) mouse model for adenine base editing detection

Base editing has emerged as a highly promising method for precise modification of the genome without a double-stranded break^16–18^. These tools comprise a D10A Cas9 nickase fused to a cytosine deaminase (C-base editor; CBE) or adenine deaminase (A-base editor; ABE) which can install a C->T or A->G change, respectively. ABEs are particularly useful for correction of human disease mutations as they can theoretically address nearly 50% of all pathogenic human single nucleotide variants (SNVs) and have a low rate of non-specific editing activity^18^. ABEs are particularly well suited to reversing stop gained mutations, which are typically pathogenic. To develop a reporter that allows for *in situ* detection of ABE activity (Gene Editing Reporter 10 (GER-10)), we mutated the glutamine residue at position 81 in mVenus to a stop codon via a C>T mutation on the plus strand. As shown in **Figure 6a**, this mutation can be converted back to a glutamine by an A>G edit on the minus strand, thus activating normal expression and fluorescence of mVenus. We used our embryo assay to validate the functionality of this allele by electroporating ABE8e RNP and measuring both fluorescence and editing levels in cultured blastocysts. As shown in **Figure 6b**, we observed strong mVenus fluorescence in ∼35% of GER10 embryos, with no evidence of leakiness in unedited embryos. Editing rates were determined by Sanger sequencing which confirmed the level of editing aligned with the degree of fluorescence for each independent experiment. Furthermore, we reverted the Q81X mutation via germline editing using the ABE8e RNP and the guides previously validated in our embryo assay and were able to demonstrate robust activation of mVenus in all cells and tissues without requiring secondary immunohistochemistry following fixation (**Figure 6c**), confirming the utility of this reporter for ABE delivery validation experiments. To further validate the use of this model *in vivo*, we delivered ABE ribonucleoprotein complexes together with an amphiphilic “shuttle peptide” that enables RNP uptake by airway epithelial cells in the lungs of GER-10 mice after intranasal instillation, **Figure 6d** ^27^. As shown in Figure 6c and 6f, edited cells of both the small and large airways displayed bright green fluorescent signal, and entry into these ciliated and mucus producing epithelial cells was enhanced by shuttle peptides. These data demonstrate that GER-10 can robustly report on ABE-mediated editing *in vivo*.

**Figure 6.**
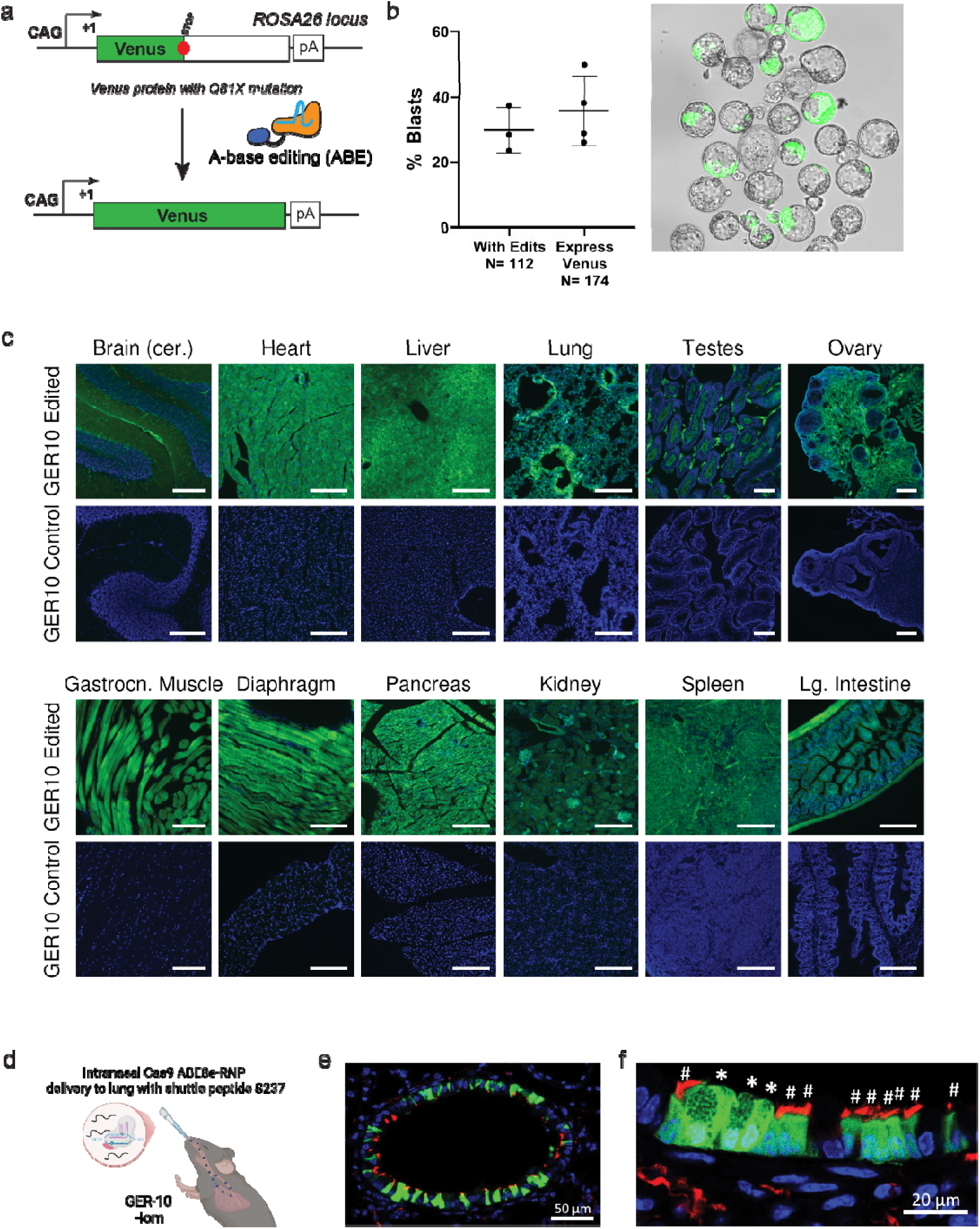
The Gene Editing Reporter 10 (GER-10) model detects A-base editing *in vivo*. **a)** Schematic of the GER-10 reporter allele. A C>T transition at position 241 results in a Q81X mutation that disrupts the expression of mVenus. A-base editing on the minus strand will reverse the mutation resulting in proper expression of the reporter. **b)** Mouse blastocyst assay showing ∼35% editing following zygote electroporation with ABE8e RNPs, with a strong correspondence between fluorescence and editing. Plots are showing mean and standard deviation of replicates with total number of blasts assessed across all replicates below each bar. **c)** Expression analysis in tissues from germline edited animals showing robust, ubiquitous expression of the reporter. Scale bar = 200 microns. d) Example of *in vivo* editing of airway epithelial cells following delivery of ABE RNPs with a shuttle peptide to the lungs. Confocal microscopy of mouse lungs showing both small (e) and large airways (f), with asterisks denoting mucus producing goblet cells, and pound signs denoting ciliated cells. Green – Venus expressing edited cells, red – ciliated cells stained with acetyl-alpha-tubulin antibody.

## Discussion

Here we describe three new mouse reporter alleles that extend and complement the growing repertoire of mouse strains available for *in vivo* assessment of genome editing activity. These models offer several key advantages. First, both the TLR-2 and TLR-7 models provide a means to monitor both HDR and NHEJ-mediated indel formation in the same allele, allowing for direct comparison of rates of each type of repair outcome. This could be useful for the optimization of HDR events, particularly where indel formation is less desirable. Second, our TLR-7 model facilitates the use of an ssODN donor, a common approach to HDR that is not feasible using the TLR-2 model. TLR-7 also includes an improved red fluorescent cassette (Katushka2S) that has enhanced spectral qualities over TagRFP. Finally, our GER-10 model provides a robust readout for A-base editing activity, a promising and therapeutically relevant genome editing technology for which limited *in vivo* resources exist.

Although our results clearly demonstrate the utility of our new models, we note a few limitations that could inform on the future development of improved models. For example, similar to Ai9, we do not observe 1:1 correspondence between editing and reporter activation as only some edits shift TagRFP or Katushka2S into the correct reading frame. By screening multiple guide options, we were able to identify guide options that show bias for the desired frame, with approximately 55% of edits producing a +2 frameshift. While similar to the edit to activation ratio seen for Ai9 (^12^ and unpublished data), there is considerable room for improvement. We are currently exploring designs that incorporate targets that are predicted to result in highly biased repair for next-generation reporters. Furthermore, our SauCas9 guide RNA targets in TLR-2 are less efficient and poorly biased towards repair into the +2 frame, resulting in relatively low activation overall. Given the common use of this Cas9 variant for in vivo delivery, future models should incorporate validated SauCas9 guide targets into the design.

The growth in the number of new editor platforms presents new opportunities and challenges to develop matched reporter tools. Indeed several new reporter models with various functionality have been recently reported ^28–30^, which complement the alleles described here and those developed as part of the SCGE program (^24^; RRID:MMRRC_068227-JAX; MMRRC_071742-JAX). While all three of our platforms are theoretically useful for detection of prime editing^15^ events, other editing modalities will require the development of new alleles. For example, given the development of new C-base editor (CBE) deaminases with improved specificity^31,32^, there is increasing interest in these editors for therapeutics. A CBE reporter with similar qualities to our GER-10 model would be of great interest.

Similarly, CRISPRa/i systems would benefit from technology-specific reporter models^33^. In all cases, reporters that include the broadest array of PAM sequences to accommodate the expanding repertoire of editor platforms is highly desirable, although it is unlikely a “one size fits all” tool could be developed given the diversity of PAM options and constraints on editing activity. Thus, there will be a need for ongoing development of new reporters (or modifications of existing reporters) to support the expanding catalogue of new editors and editing applications.

## Methods

### Mice

All animal studies were performed in accordance with the Guide to the Care and Use of Laboratory Animals and were reviewed and approved by the Institutional Animal Care and Use Committee of The Jackson Laboratory.

### Mouse model engineering

<u>TLR-2</u> mice (B6.129S6-*Gt(ROSA)26Sor^tm^*^12^(CAG–Venus*,–TagRFP*)*^Rkuhn^*/MurrJ (JAXStrain # 034033; RRID:IMSR_JAX:034033) were generated with a R26-GFP_KI-TLR2 (“traffic light reporter”) targeting vector was designed to insert the TLR reporter construct (“TLR-2”) into the *Gt(ROSA)26Sor* locus: the TLR reporter construct (“TLR-2”) containing a CAG promoter Venus*-P2A-TagRFP-bovine hGH polyA, in a reading frame that is shifted by 2 bp (+3) ^19^. The Venus* used here is a non-functional Venus having codons 117-152 replaced with 52bp of mouse *Gt(ROSA)26Sor* and 56bp of mouse *Rab38* (RAB38, member RAS oncogene family sequence). The targeting construct was electroporated into (C57BL/6J x 129S6/SvEvTac)F1-derived IDG3.2 embryonic stem (ES) cells. Correctly targeted ES cells were injected into blastocysts. The resulting chimeric animals were tested for germline transmission. The mice were then backcrossed to C57BL/6J for 3 generations

### TLR-7

(C57BL/6J-*Gt(ROSA)26Sor^em^*^8^(CAG–Venus*,–Katushka2S)*^Murr^*/MurrMmjax, MMRRC Strain #069711-JAX; RRID:MMRRC_069711-JAX) was developed by inserting the following construct into the second intron of the *Gt(ROSA)26Sor* locus of C57BL/6J via Bxb1 mediated integration at landing sites engineered into strain C57BL/6J-*Gt(ROSA)26Sor^em7Mvw^*/Mvw (JAXStrain #036152): CMV Enhancer, Chicken Beta-actin promoter and chimeric intron, followed by mVenus (GFP) coding sequence interrupted by the insertion of 30 bases and deletion of sequence codons 95-97, and P2A peptide.

Katushka2S (RFP) follows the P2A peptide in the +3 reading frame and bovine growth hormone polyA termination sequence. The targeting construct was microinjected into oocytes from C57BL/6J-*Gt(ROSA)26Sor^em7Mvw^*/MvwJ, along with Bxb1 integrase mRNA to generate founders with properly targeted sequences to the Rosa26 locus^34^. Mice were screened via PCR and Sanger sequencing of the targeted region to confirm proper insertion, then backcrossed to C57BL/6J for at least 2 generations and then intercrossed to generate homozygotes.

<u>GER-10</u> reporter knock-in mice (C57BL/6N-*Gt(ROSA)26Sor^em^*^3^(CAG–Venus*)*^Rkuhn^*/MurrMmjax; MMRRC Strain #069724-JAX; RRID:MMRRC_069724-JAX) were created using CRISPR/cas9 genome editing to insert a CAG promoter controlling expression of a non-functional mVenus (GFP) reporter (with a stop engineered at codon 81) in the endogenous *Gt(ROSA)26Sor* locus. The reporter construct contains a CAG promoter, mutated Venus*, bovine hGH, and polyA sequence. The targeting construct was microinjected into zygotes of C57BL/6N mice together with *cas9* protein and a guide RNA creating a DNA double strand break in the Rosa26 locus, then transferred to pseudopregnant females. Progeny were screened by DNA sequencing of the targeted region and a founder was identified harboring the desired allele. This founder was bred to C57BL/6N for germline transmission. The colony was backcrossed to C57BL/6N for at least two generations.

Annotated reporter cassette sequences are provided in the Supplementary Material.

### In vitro mouse embryo assay

#### Guide RNA annealing and ribonucleoprotein (RNP) complex formation

Model specific crRNA is annealed with trRNA following IDT AltR System protocols. Briefly, both components are resuspended at 100 μM in IDT Duplex Buffer, combined in equal amounts, heated to 95°C for 5 mins, and allowed to cool passively to room temperature. Following this annealing step, concentration of annealed guide RNA is assayed by NanoDrop. Crispr:tracr guide RNA hybrid is complexed with AltR Cas9 at 37°C for 15 mins in a thermocycler.

#### Reporter Testing via RNP Electroporation and blastocyst culture

Reporter gRNAs are individually tested *ex vivo* to assess reporter efficacy. Super-ovulated C57BL/6NJ or C57BL/6J females were mated to reporter male studs, and zygotes were harvested and placed in a droplet of 10 μl TE with AltR-Cas9 (500 ng/μl), guide(s) RNA (600 ng/μl) preassembled as described above, and if used, ssDNA repair template donor (2000 ng/μl) These components were combined with 10 μl low serum media (Opti-Mem, Sigma-Millipore), and transferred to an electroporation cuvette (Harvard Apparatus) with a 1mm gap electrode. Using a BTX ECM830 Electro Square Porator (Harvard Apparatus), embryos were electroporated with six 3 ms pulses of 30V separated by 100 ms each ^35^. Following CRISPR/Cas9 electroporation, zygotes are cultured in Sydney Cleavage Medium (COOK Medical) at 37°C under CO_2_ in a benchtop incubator (COOK Medical). After 96 hours, blastocysts are collected for imaging and DNA extraction. Blastocysts are imaged on a Leica Dmi8 inverted fluorescent microscope at 10X magnification with an NA0.3 Ph1 objective, DFC9000GT sCMOS camera using the LasX Navigator acquisition software. For DNA extraction, blastocysts are collected in individual PCR tubes with 1.5 μl 25 mM NaOH / 0.2 mM EDTA at 95°C for 15 minutes and neutralized with an equal volume of 40 mM Tris HCl. The genomic region of interest is PCR amplified from 3 μl of DNA template, and product was Sanger Sequenced to assess CRISPR editing outcomes (Primer sequences provided in **Supplementary Table S2**). Sanger sequence traces were analyzed using the ICE (Inference of CRISPR Editing) deconvolution tool from Synthego. (https://ice.synthego.com; Synthego Performance Analysis, ICE Analysis. 2019. v2.0.)

### Germline editing for validation of widespread expression

Zygotes from C57BL/6NJ or C57BL/6J female mice mated to reporter males were harvested and electroporated with AltR-Cas9 (500 ng/μl) and guide RNA (600 ng/μl) as described above. Oviduct transfers into pseudopregnant dams were performed immediately following electroporation. Pregnancies were allowed to progress to term and offspring were genotyped by Sanger sequencing. Animals with edits predicted to activate the reporter were mated to inbred females and at least 3 edited offspring per sex and from more than one founder line were assessed.

### Tissue sampling and imaging

Tissues from each animal are collected, fixed in 4% PFA or NBF for 2-4 hours, cryoprotected in 30% sucrose and embedded in OCT medium and frozen on dry ice. Tissues are cryosectioned and either directly imaged (GER-10) or incubated with primary antibodies specific to the fluorophore and corresponding secondary antibodies (TLR-2, TLR-7) before staining with DAPI/Hoechst and imaged.

Images are assessed for reporter activation potential across tissues. TagRFP and Katushka2S proteins are detected using rabbit anti-RFP polyclonal antibody (Invitrogen Cat # R10367) and corresponding secondary antibodies such as Cy3-Donkey anti-Rabbit (Jackson ImmunoResearch #711-165-52). A standard panel of tissues is collected and assessed for each reporter line including brain, eye, gastrocnemius muscle, heart, lung, diaphragm, skin, thymus, lymph node, kidney, spleen, liver, stomach, small intestine, large intestine, adrenal gland, pancreas, bladder, ovary/testes, uterus/epididymis. Images were collected using Leica Dmi8 inverted fluorescent microscope at 10x magnification, NA 0.3 Ph1 objective, DFC9000GT sCMOS camera and LasX Navigator acquisition software. Fluorochromes were detected using Chroma filter cubes with the following excitation (ex.) and emission (em.) properties: DAPI (ex. 404 nm, em.420-450 nm), GFP (ex. 475 nm, em. 506-532 nm), RFP (ex. 555 nm, em. 578-610 nm). Fluorescent images were rendered using OMERO.figure with RFP and GFP baseline thresholds set using untreated or unedited littermate or colony controls.

### In vivo delivery of AAV and LNP

#### AAV delivery of SauCas9 and guide

Adult homozygous TLR-2 mice (10 weeks of age) were administered intravenously via tail vein, 1 x 10^11^vg of custom synthesized AAV9 vector (Vigene) encoding both SauCas9 nuclease and SaCa9-g3. Age matched control TLR-2 mice received saline. Mice were euthanized 4 weeks later and tissues collected for both sequencing and imaging as described above. Quantification of editing activity was performed by analyzing the Sanger sequence traces with the Synthego ICE evaluation tool.

#### Delivery of ABE8e-RNP +S237 peptide to mouse lung, staining and confocal analysis

Adult GER-10 mice (8-10 weeks of age) were lightly anesthetized and instilled intranasally with 2 doses of ABE8e RNP [ABE8e-Cas9 (5μM)/sgRNA (4 μM)] +shuttle peptide S237 (80μM), 24 hours apart, and tissues harvested 8 days after the last instillation. Mice were euthanized and vasculature perfused via the heart with ice cold PBS, followed by cold 4% PFA. Trachea, bronchi and lungs were fixed for 4 hours in 4% PFA, equilibrated in 30% sucrose/PBS and shipped to the processing laboratory (U Iowa). Lung tissues were then embedded in OCT and stored at -80°C prior to sectioning. Lung tissues were sectioned at 10μm for imaging. Ciliated cells in large and small airways were stained with acetyl-alpha-tubulin antibodies (Cell Signaling Technology, Ref #5335, and Invitrogen, Ref #A-21244, goat anti-rabbit secondary antibody – Alexa Fluor 647), edited cells express mVenus green fluorescent protein, and nuclei were stained with DAPI. Confocal microscopy images were collected with Zeiss LSM 710 confocal microscope equipped with Zen2009 software. Image composites were prepared using ImageJ-FIJI (NIH).

## Statistical Assessment

Data are presented as means of technical replicates with standard deviation and described in the legends for each figure and no data were excluded. Samples sizes were not predetermined using statistical methods. Allocation of mice into groups was performed independently and randomly based on availability of animals. Both males and female were included in each study design and roughly equal numbers of each sex were included for each.

## Data Availability

Mouse strains associated with this publication are available through the Mutant Mouse Regional Resource Centers or the Jackson Laboratory Repository (RRID:IMSR_JAX:034033, RRID:MMRRC_069711-JAX, RRID:MMRRC_069724-JAX). Plasmids associated with model creation or repair have been deposited to Addgene (https://www.addgene.org) under Addgene IDs 64322, 64215. Source data are provided with this publication.

## Acknowledgements

We thank other members of the SCGE consortium including members of the animal model working group and NIH Program staff for their thoughts and feedback on the generation and characterization of these models. We thank the expert assistance and support from The Jackson Laboratory Imaging Sciences, Genome Technologies, and Genetic Engineering Technology Cores which are supported in part by the JAX Cancer Center Support Grant (P30CA034196). The content is solely the responsibility of the authors and does not necessarily represent the official views of the National Institutes of Health.

## Author contributions

Conceptualization, S.A.M., R.K., W.W., DEB; Methodology, K.J.S., R.K., W.W., S.A.M., M.V.W., D.E.B., J.R., T.D.; Formal analysis, K.J.S.; Investigation, K.J.S., E.S., C.H., Y.G., T.L.D., L.B., S.H., J.D., K.K., J.R., A.C., T.D., B.E.L.; Resources, R.K., W.W., D.G., C.M.L., S.A.M.; Data Curation, K.J.S., Y.G., R.K.; Writing (original draft) K.J.S., S.A.M.; Writing (reviewing and editing,) K.J.S., B.E.L., K.K., P.B.M., J.D., R.K., W.W., S.A.M.; Visualization, K.J.S, R.K., S.A.M.; Supervision, K.J.S., W.W., R.K., P.B.M, D.E.B.; Project Administration, K.J.S., W.W., R.K., S.A.M.; Funding Acquisition S.A.M., C.M.L., P.B.M.

