## Supplementary Information for "Novel mouse reporter models for the detection of genome editing events in vivo"

This file includes:

Supplementary Figures S1-3 and

Supplementary Tables S1 and S2

Supplementary Methods including annotated sequences of reporter allele cassettes.

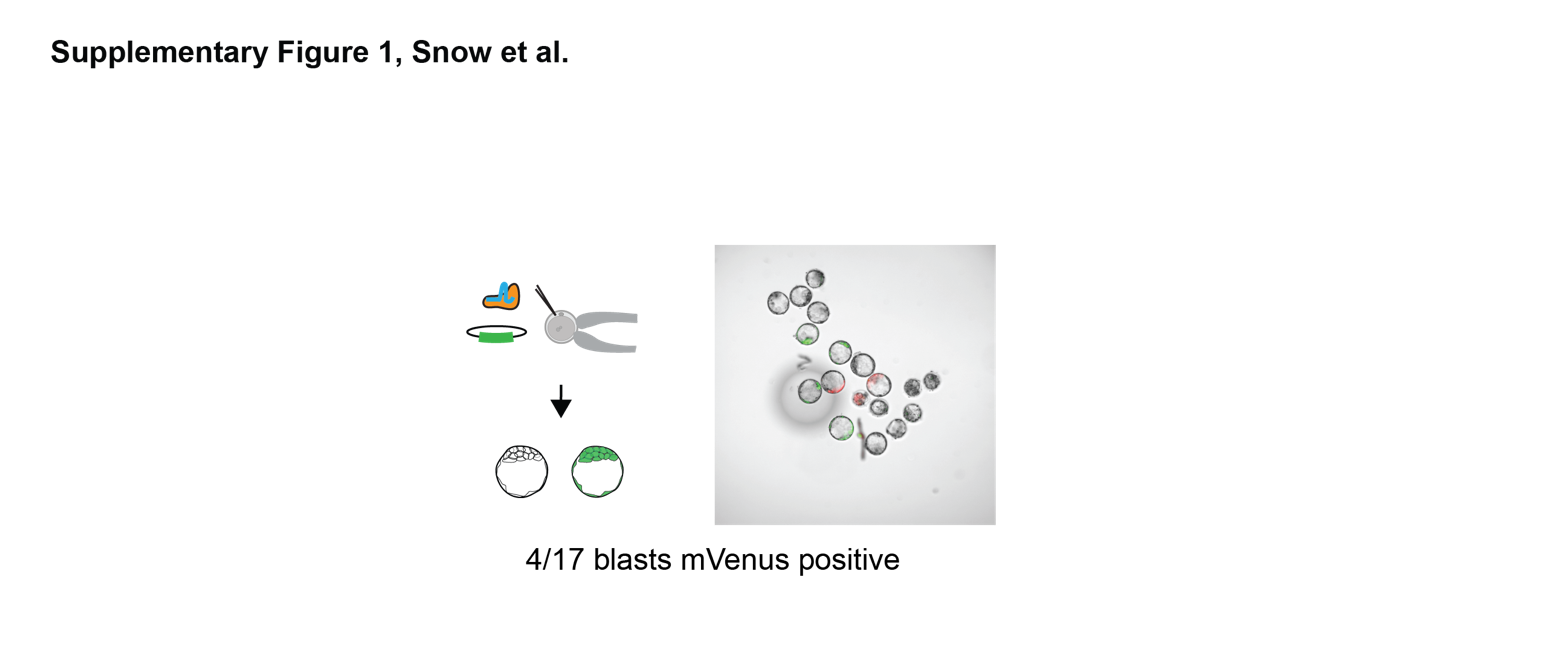

**Supplementary Figure S1**. Detection of HDR editing events using our blast editing and culture assay described in Figure 1b. Successful mVenus repair following SpyCas9 editing via electroporation along with the supplied plasmid donor template results in GFP expression in blastocysts while NHEJ resolution of dsDNA breaks with +2 indel allows for TagRFP expression.

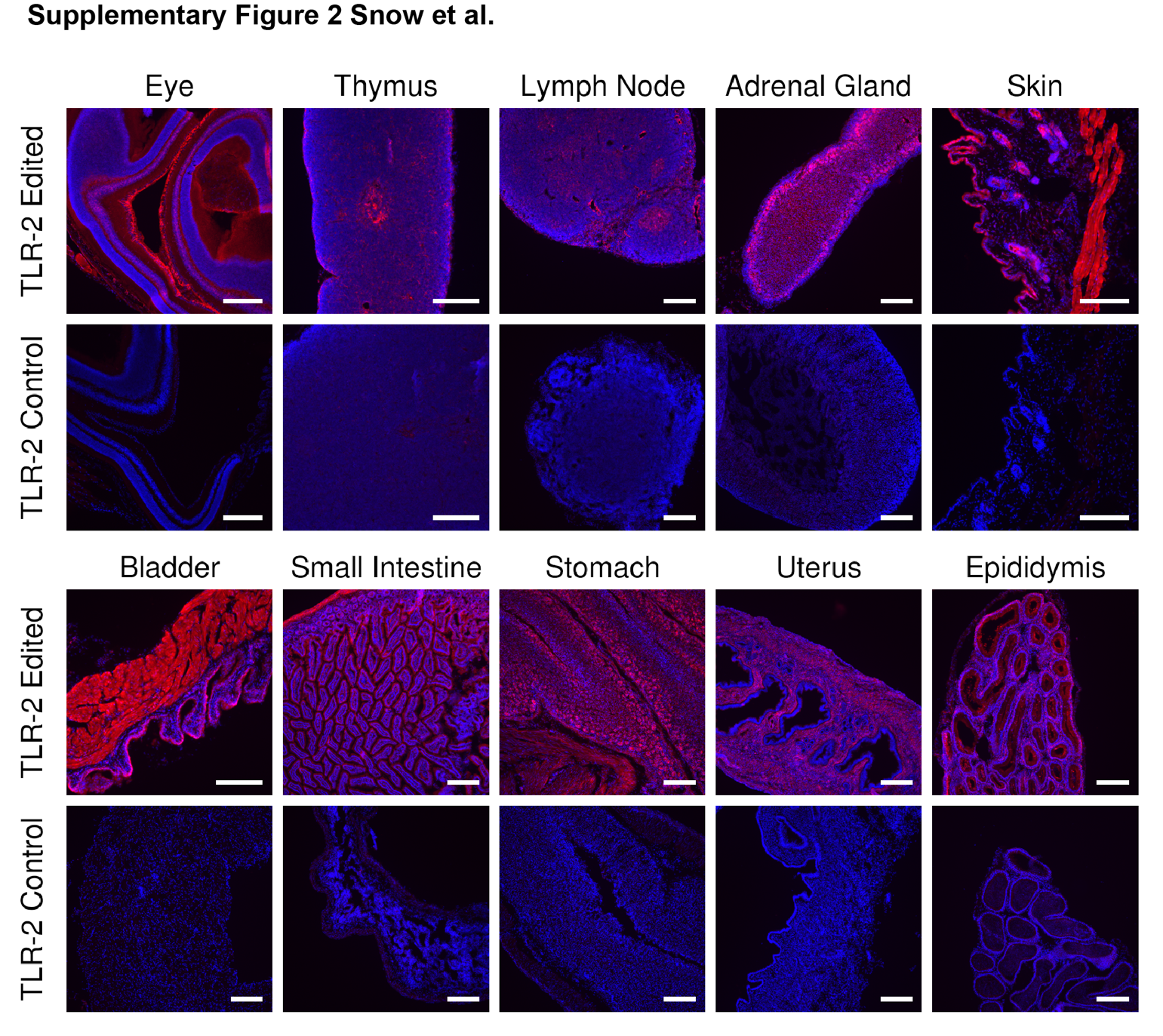

**Supplementary Figure S2**. **Widespread** **expression of TLR-2 reporter allele in various tissues**. TLR-2 mice with a germline recombined reporter allele (as described in Figure 3) display widespread reporter activation in an expanded tissue set. Scale bar = 200 microns.

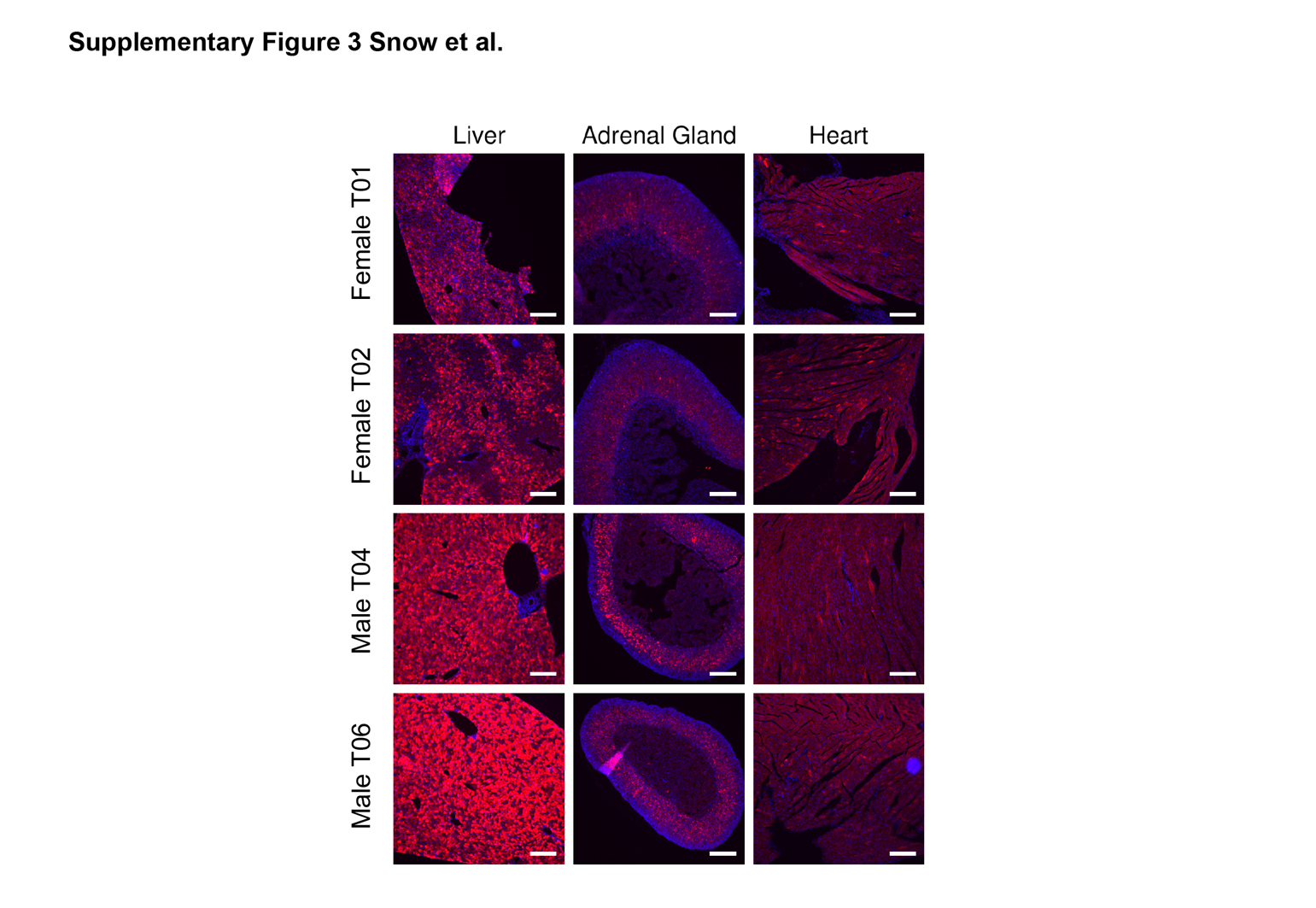

**Supplementary Figure S3**. **Intravenous delivery of AAV9-SauCas9-SaC9g3 into adult mice activates the TLR-2 reporter allele** **in tissues beyond the liver**. As described in Figure 5, AAV delivery of SauCas9 + guide edits cells beyond the liver, including the adrenal gland and heart. Representative images from 2 males and 2 females treated with the AAV (1x 10^11^vg AAV9 per mouse). Scale bars = 200 microns.

**Supplementary Tables 1&2**

**Supplementary Table 1: Guide RNA sequences used in this study**

| Reporter Model | Guide Name | Sequence | PAM | Nuclease | DNA Strand |
| --- | --- | --- | --- | --- | --- |
| TLR-2 | R26-1* | ACTCCAGTCTTTCTAGAAGA | TGG | SpyCas9 | Fwd |
|  | R26-2 | CGCCCATCTTCTAGAAAGAC | TGG | SpyCas9 | Rev |
|  | R26-3 | TTGCAGCTCGAACTTCACCT | CGG | SpyCas9 | Rev |
|  | R26-52* | GGTAGCGGGCGAAGCACTGC | AGG | SpyCas9 | Rev |
|  | R26-59* | GACGTAGCCTTCGGGCATGG | CGG | SpyCas9 | Rev |
|  | R26-69 | CAACTACAAGACCCGCGCCG | AGG | SpyCas9 | Fwd |
|  | SaC9-g1 | AAGCACTGCAGGCCGTAGCC | CAGGGT | SauCas9 | Rev |
|  | SaC9-g2 | GTCGTGCTGCTTCATGTGGT | CGGGGT | SauCas9 | Rev |
|  | SaC9-g3 | AGCTCGAACTTCACCTCGGC | GCGGGT | SauCas9 | Rev |
| TLR-7 | TLR7-g2-Rev | ACTCCATCTTCTAGAAAGAC | TGG | SpyCas9 | Rev |
| GER10 | GER10-ABE-REV | CGTGCTACTTCATGTGGTCG | GGG | ABE8e (SpyCas9) | Rev |
| * Compatible with TLR-7 | | | | | |

**Supplementary Table 2: Genotyping primers for editing assessment**

| Reporter Model | Primer Name | Sequence | Expected Product Size (unedited) |
| --- | --- | --- | --- |
| TLR-2 | 4015 Venus F1 | AGCTGACCCTGAAGCTGAT |  |
|  | 4016 Venus R1 | GACGTTGTGGCTGTTGTAGTT | 393 |
|  | 45714 Venus F | CAGAAGAACGGCATCAAGG |  |
|  | 45716 TagRFP R | TCAGCTCTTCGCCCTTAGAC | 339 |
| TLR-7 | 4015 Venus F1 | AGCTGACCCTGAAGCTGAT |  |
|  | 4016 Venus R1 | GACGTTGTGGCTGTTGTAGTT | 367 |
| GER10 | 4015 Venus F1 | AGCTGACCCTGAAGCTGAT |  |
|  | 4016 Venus R1 | GACGTTGTGGCTGTTGTAGTT | 329 |
| Blastocyst Primers | TLR-pCAG-GT-F | TCTGCTAACCATGTTCATGCCT |  |
|  | TLR-pCAG-GT-R | TCTCGTTGGGGTCTTTGCTC | ~1kb |

**Supplementary Method: Annotated reporter cassette sequences**

**TLR-2 Reporter sequence**

Venus coding region (frame 1)

108 bp from Rosa26 and Rab38 locus replacing Venus codons 117 – 152, **TAG** stop codon in red

*R26-2* guide sequence in *italics*, underlined and shaded, PAM sequence lowercase and underlined (reverse strand)

*R26-52* guide sequence in *italics*, underlined and shaded, PAM sequence lowercase and underlined (reverse strand)

(see Supplementary Table 1 for additional guide sequences)

Spacer 8 bp

P2A peptide (frame +3)

TagRFP (frame +3)

ATGGTGAGCAAGGGCGAGGAGCTGTTCACCGGGGTGGTGCCCATCCTGGTCGAGCTGGACGGCGACGTAAACGGCCACAAGTTCAGCGTGTCCGGCGAGGGCGAGGGCGATGCCACCTACGGCAAGCTGACCCTGAAGCTGATCTGCACCACCGGCAAGCTGCCCGTGCCCTGGCCCACCCTCGTGACCACCCTGGGCTACGGcct*GCAGTGCTTCGCCCGCTACC*CCGACCACATGAAGCAGCACGACTTCTTCAAGTCCGCCATGCCCGAAGGCTACGTCCAGGAGCGCACCATCTTCTTCAAGGACGACGGCAACTACAAGACCCGCGCCGAGGTGAAGTTCGAGCTGCAACTcca*GTCTTTC****TAG****AAGATGGGCG*GGAGTCTTCTGGGCAGGCTTATATCAAGCGCTATGTGCACCAAAACTTCTCCTCGCACTACCGGGCCACCATTGGTGATCACCGCCGACAAGCAGAAGAACGGCATCAAGGCCAACTTCAAGATCCGCCACAACATCGAGGACGGCGGCGTGCAGCTCGCCGACCACTACCAGCAGAACACCCCCATCGGCGACGGCCCCGTGCTGCTGCCCGACAACCACTACCTGAGCTACCAGTCCGCCCTGAGCAAAGACCCCAACGAGAAGCGCGATCACATGGTCCTGCTGGAGTTCGTGACCGCCGCCGGGATCACTCTCGGCATGGACGAGCTGTACAAGTAGACGCGTTGGCCACGAACTTCTCTCTGTTAAAGCAAGCAGGAGATGTTGAAGAAAACCCCGGGCCTATGGTGTCTAAGGGCGAAGAGCTGATTAAGGAGAACATGCACATGAAGCTGTACATGGAGGGCACCGTGAACAACCACCACTTCAAGTGCACATCCGAGGGCGAAGGCAAGCCCTACGAGGGCACCCAGACCATGAGAATCAAGGTGGTCGAGGGCGGCCCTCTCCCCTTCGCCTTCGACATCCTGGCTACCAGCTTCATGTACGGCAGCAGAACCTTCATCAACCACACCCAGGGCATCCCCGACTTCTTTAAGCAGTCCTTCCCTGAGGGCTTCACATGGGAGAGAGTCACCACATACGAAGACGGGGGCGTGCTGACCGCTACCCAGGACACCAGCCTCCAGGACGGCTGCCTCATCTACAACGTCAAGATCAGAGGGGTGAACTTCCCATCCAACGGCCCTGTGATGCAGAAGAAAACACTCGGCTGGGAGGCCAACACCGAGATGCTGTACCCCGCTGACGGCGGCCTGGAAGGCAGAAGCGACATGGCCCTGAAGCTCGTGGGCGGGGGCCACCTGATCTGCAACTTCAAGACCACATACAGATCCAAGAAACCCGCTAAGAACCTCAAGATGCCCGGCGTCTACTATGTGGACCACAGACTGGAAAGAATCAAGGAGGCCGACAAAGAGACCTACGTCGAGCAGCACGAGGTGGCTGTGGCCAGATACTGCGACCTCCCTAGCAAACTGGGGCACAAACTTAATTGA

**TLR-7 Reporter Sequence**

mVenus coding region (frame 1)

Spacer sequence (30 bp) replacing codons 96-98, **TAG** stop codon in red.

*TLR7-g2-REV*, guide sequence in *italics* and shaded, PAM sequence lowercase and underlined (reverse strand)

*R26-52* guide sequence in *italics*, underlined and shaded, PAM sequence lowercase and underlined (reverse strand)

(see Supplementary Table 1 for additional guide sequences)

Spacer (1bp)

P2A peptide (frame 3)

Katushka2S (frame 3)

ATGGTGAGCAAGGGCGAGGAGCTGTTCACCGGGGTGGTGCCCATCCTGGTCGAGCTGGACGGCGACGTAAACGGCCACAAGTTCAGCGTGTCCGGCGAGGGCGAGGGCGATGCCACCTACGGCAAGCTGACCCTGAAGCTGATCTGCACCACCGGCAAGCTGCCCGTGCCCTGGCCCACCCTCGTGACCACCCTGGGCTACGGcct*GCAGTGCTTCGCCCGCTACC*CCGACCACATGAAGCAGCACGACTTCTTCAAGTCCGCCATGCCCGAAGGCTACGTCCAGACTcca*GTCTTTC****TAG****AAGATGGAGT*ACCCATCTTCTTCAAGGACGACGGCAACTACAAGACCCGCGCCGAGGTGAAGTTCGAGGGCGACACCCTGGTCAACCGCATCGAGCTCAAGGGCATCGACTTCAAGGAGGACGGCAACATCCTGGGGCACAAGCTGGAGTACAACTACAACAGCCACAACGTCTATATCACCGCCGACAAGCAGAAGAACGGCATCAAGGCCAACTTCAAGATCCGCCACAACATCGAGGACGGCGGCGTGCAGCTCGCCGACCACTACCAGCAGAACACCCCCATCGGCGACGGCCCCGTGCTGCTGCCCGACAACCACTACCTCAGCTACCAGTCCGCCCTCAGCAAAGACCCCAACGAGAAGCGCGATCACATGGTCCTGCTGGAGTTCGTCACCGCCGCCGGGATCACTCTCGGCATGGACGAGCTGTACAAGTAGCCGCCACGAACTTCTCTCTGTTAAAGCAAGCAGGAGATGTTGAAGAAAACCCCGGGCCTATGGTGGGTGAGGATAGCGTGCTGATCACCGAGAACATGCACATGAAACTGTACATGGAGGGCACCGTGAACGACCACCACTTCAAGTGCACATCCGAGGGCGAAGGCAAGCCCTACGAGGGCACCCAGACCATGAAGATCAAGGTGGTCGAGGGCGGCCCTCTCCCCTTCGCCTTCGACATCCTGGCTACCAGCTTCATGTACGGCAGCAAAACCTTTATCAACCACACCCAGGGCATCCCCGACTTCTTTAAGCAGTCCTTCCCTGAGGGCTTCACATGGGAGAGGATCACCACATACGAAGACGGGGGCGTGCTGACCGCTACCCAGGACACCAGCCTCCAGAACGGCTGCCTCATCTACAACGTCAAGATCAACGGGGTGAACTTCCCATCCAACGGCCCTGTGATGCAGAAGAAAACACTCGGCTGGGAGGCCAGCACCGAGATGCTGTACCCCGCTGACAGCGGCCTGAGAGGCCATGCCCAGATGGCCCTGAAGCTCGTGGGCGGGGGCTACCTGCACTGCTCCCTCAAGACCACATACAGATCCAAGAAACCCGCTAAGAACCTCAAGATGCCCGGCTTCTACTTCGTGGACCGGAGACTGGAAAGAATCAAGGAGGCCGACAAAGAGACCTACGTCGAGCAGCACGAGATGGCTGTGGCCAGATACTGCGACCTGCCTAGCAAACTGGGGCACAGCTAA

**TLR-7 Reporter mVenus repair sequences**

*9 bp repair sequence in Venus coding region underlined below (codon 96-98)*

131 bp asymmetrical ssODN repair sequence:

GGGCTACGGCCTGCAGTGCTTCGCCCGCTACCCCGACCACATGAAGCAGCACGACTTCTTCAAGTCCGCCATGCCCGAAGGCTACGTCCAG**GAGCGCACC**ATCTTCTTCAAGGACGACGGCAACTACAAGA

200 bp symmetrical ssODN repair sequence:

CCCTGGGCTACGGCCTGCAGTGCTTCGCCCGCTACCCCGACCACATGAAGCAGCACGACTTCTTCAAGTCCGCCATGCCCGAAGGCTACGTCCAG**GAGCGCACC**ATCTTCTTCAAGGACGACGGCAACTACAAGACCCGCGCCGAGGTCAAGTTCGAGGGCGACACCCTGGTCAACCGCATCGAGCTCAAGGGCATCGA

*131 bp asymmetrical ssODN highlighted in yellow*

**GER-10 Reporter**

mVenus coding sequence with target Q81X (tAG) stop codon in red

GER10-ABE-REV guide sequence in *italics* and shaded, PAM sequence lowercase and underlined (reverse strand)

ATGGTGAGCAAGGGCGAGGAGCTGTTCACCGGGGTGGTGCCCATCCTGGTCGAGCTGGACGGCGACGTAAACGGCCACAAGTTCAGCGTGTCCGGCGAGGGCGAGGGCGATGCCACCTACGGCAAGCTGACCCTGAAGCTGATCTGCACCACCGGCAAGCTGCCCGTGCCCTGGCCCACCCTCGTGACCACCCTGGGCTACGGCCTGCAGTGCTTCGCCCGCTAccc*CGACCACATGAAG****tAG****CACG*ACTTCTTCAAGTCCGCCATGCCCGAAGGCTACGTCCAGGAGCGCACCATCTTCTTCAAGGACGACGGCAACTACAAGACCCGCGCCGAGGTGAAGTTCGAGGGCGACACCCTGGTGAACCGCATCGAGCTGAAGGGCATCGACTTCAAGGAGGACGGCAACATCCTGGGGCACAAGCTGGAGTACAACTACAACAGCCACAACGTCTATATCACCGCCGACAAGCAGAAGAACGGCATCAAGGCCAACTTCAAGATCCGCCACAACATCGAGGACGGCGGCGTGCAGCTCGCCGACCACTACCAGCAGAACACCCCCATCGGCGACGGCCCCGTGCTGCTGCCCGACAACCACTACCTGAGCTACCAGTCCGCCCTGAGCAAAGACCCCAACGAGAAGCGCGATCACATGGTCCTGCTGGAGTTCGTGACCGCCGCCGGGATCACTCTCGGCATGGACGAGCTGTACAAGTAA

Guide sequence for A-base editing on reverse strand (5’-3’), target adenine in orange: CGTGCT**a**CTTCATGTGGTCG GGG
